# Machine Learning Classification Reveals Robust Morphometric Biomarker of Glial and Neuronal Arbors

**DOI:** 10.1101/2022.04.02.486839

**Authors:** Masood A. Akram, Qi Wei, Giorgio A. Ascoli

## Abstract

Neurons and glia are the two main cell classes in the nervous systems of most animals. Although functionally distinct, neurons and glia are both characterized by multiple branching arbors stemming from the cell bodies. Glial processes are generally known to form smaller trees than neuronal dendrites. However, the full extent of morphological differences between neurons and glia in multiple species and brain regions has not yet been characterized, nor is it known whether these cells can be reliably distinguished based on geometric features alone. Here, we show that multiple supervised learning algorithms (K-nearest neighbor, random forest, and support vector machine) deployed on a large database of morphological reconstructions can systematically classify neuronal and glial arbors with nearly perfect accuracy and precision. Moreover, we report multiple morphometric properties, both size-related and size-independent, that differ substantially between these cell types. In particular, we newly identify an individual morphometric measurement, Average Branch Euclidean Length (ABEL) that can robustly separate neurons from glia across multiple animal models, a broad diversity of experimental conditions, and anatomical areas, with the notable exception of the cerebellum. We discuss the practical utility and physiological interpretation of this discovery.

## Introduction

Neuronal classification is an increasingly important subject because of its ultimate goal of linking cell types with computation, behavior, and cognition (Armañanzas & Ascoli, 2015). The main experimental approaches to characterize neurons are biochemistry, physiology, and morphology (Petilla Interneuron Nomenclature Group, 2008). These techniques have all yielded major breakthroughs in recent years thanks to rapid progress in genomics and transcriptomics, large-scale electric recordings, and high-resolution microscopic imaging (Litvina et al., 2019), respectively. Both the European Human Brain Project and the American BRAIN Initiative identified cell type classification among their first priorities (Insel et al., 2013; Markram, 2012). Relative to neurons, glial cells have received less attention despite being similarly abundant in most organisms with a nervous system, including humans and all common animal models. Glia are involved in numerous important functions, such as myelination, antiinflammatory protection, maintenance of neurochemical environment, and exchanges between nervous and vascular systems (Aguzzi et al., 2013; Bronzuoli et al., 2018; Jessen, 2004; Rasband, 2016). Most glial cells emanate from the cell body complex branching processes that resemble the structural architecture of neuronal dendrites. While large numbers of neurons have been morphologically reconstructed for over three decades, digitally tracing glial trees has only more recently become a routine practice as well.

Although it is usually recognized that glial arbors are smaller than dendritic trees (García-Marín et al., 2007; Lu et al., 2015; Veldman et al., 2020; Zisis et al., 2021), a comprehensive morphological comparison has not yet been carried out. In particular, it is still unknown whether these two main categories of cells can be reliably distinguished based on geometric features alone. The general problem is further complicated by several factors. First, both neurons and glia are intrinsically diverse, with the former often distinguished by circuit role (long-range projecting, local interneurons, and sensory receptors) and the latter typically divided by functional specialization (microglia, astrocytes, oligodendrocytes, etc.). Second, both neurons and glia tend to differ broadly across animal species (especially between vertebrates and invertebrates), anatomical regions (e.g., neocortex, brainstem, spinal cord, peripheral nervous system), and developmental stage (such as embryo, early postnatal, and adult). Third, morphological characterization may be affected by the tremendous variability in experimental methods, including animal care, histological details, labeling protocol, imaging modality, and reconstruction software. Thus, it remains an open question whether suitable morphometric biomarkers exist that can robustly and systematically discriminate between neuronal and glial arbors.

Hundreds of laboratories worldwide continuously contribute their digital reconstructions of neurons and glia to the public online database NeuroMorpho.Org (Akram et al., 2018). This repository associates every cell entry with metadata (Bijari et al., 2020) describing the animal subject (species, strain, sex, age, and weight), anatomy (brain region, sub-region, cell type, and sub-type), experimental details (protocol, condition, histology, microscopy, and tracing), and provenance (authors, source publication, original version, and processing logs). Moreover, the detailed 3D representation of arbor geometry is accompanied by a battery of morphometric parameters extracted with L-Measure (Scorcioni et al., 2008), such as total length, number of branches, arbor height, and tortuosity. Glial cells were introduced to NeuroMorpho.Org in version 7.1 (2017) and now constitute 11.3% of over 170,000 tracings. The unrestricted availability of these data provides an unprecedented opportunity for scientific exploration, statistical analysis, and computational modeling (Ascoli et al., 2017).

Machine learning is a branch of artificial intelligence that aims at enabling the efficient and automatic detection of data patterns. Machine learning strives to produce the most accurate predictions, a distinct goal from that of statistical models designed to quantify the relationships variables (Aha et al., 1991). Recent advancements in machine learning have not only benefitted healthcare with automatic diagnoses and treatment planning (Kohli & Arora, 2018), but were also successfully applied in neurobiological data analysis such as automatic tracing of neurons and glia (Peng et al., 2017) and their quantification (Bijari et al., 2021). In supervised machine learning, an algorithm is trained with the known class labels and identifies the most informative combination of features that are associated with those labels (Kotsiantis et al., 2006). The resultant classifiers can then be applied for predicting the labels of unknown data based on their feature values. Here we leverage supervised learning algorithms (K-Nearest Neighbor, Random Forest, and Support Vector Machine) to classify glia and neurons, and to recognize the morphological structures that distinguish these two main cell types of the nervous system.

## Materials and Methods

### Dataset selection and preprocessing

Morphological reconstructions of neurons and glia were obtained from NeuroMorpho.Org using the *Summary Reporting* web-based functionality (Akram et al., 2022). This tool collates for every digital tracings the annotation of 35 distinct metadata fields, providing a detailed qualitative description of the cell (Parekh et al., 2015), as well as 21 morphometric measurements which capture the quantitative structural features of the arbor (Scorcioni et al., 2008). First, we downloaded all glial cells available at the time we began data analysis (Fall 2020), corresponding to 10 consecutive releases of NeuroMorpho.Org (versions 7.1 to 8.0 inclusive). We then analyzed the distributions of their metadata with respect to animal species, developmental stage, anatomical region, and other experimental conditions, and queried the database to identify the same number of neurons with the most similar metadata characteristics. Since we were interested in comparing glial processes specifically to neuronal dendrites (as opposed to axons), only neurons with dendritic tracings available were selected. Moreover, we solely included neurons and glia with complete or moderately complete reconstructions, thus excluding those annotated as incomplete dendrites or incomplete glial processes by the original contributors. The resulting balanced dataset of 22,792 cells was comprised of 11,398 neurons and 11,394 glia.

Of the 21 morphometrics extracted for each cell from NeuroMorpho.Org, we excluded Soma Surface and Depth from the analysis. Soma Surface is not an arbor morphometric, and 4.7% of the tracings in our dataset did not include soma reconstruction. Depth was similarly not reported for 8.6% of the cells as the accuracy of the tracing is reduced in certain cases by light diffraction and tissue shrinkage in the direction perpendicular to the imaging place. The remaining 19 morphometric features were used in the analysis: number of stems, number of bifurcations, number of branches, overall width, overall height, average diameter, total length, total surface, total volume, maximum Euclidean distance, maximum path distance, maximum branch order, average contraction, total fragmentation, partition asymmetry, average Rall’s ratio, average bifurcation angle local, average bifurcation angle remote, and fractal dimension. The formal definitions of these metrics are available on the Frequently Asked Questions of NeuroMorpho.Org (http://neuromorpho.org/myfaq.jsp) and on the online manual of L-Measure (http://cng.gmu.edu:8080/Lm/help/index.htm).

Most reconstructions in NeuroMorpho.Org have coordinates reported in microns. In a subset of reconstructions, however, the coordinates are expressed in pixels. In these cases, the nominal measurements listed in the morphometric tables must be converted by an appropriate scaling factor. Therefore, we manually calculated the height of at least one cell in each archive from the figures of the corresponding original publications (and relative scale bar) and compared the resulting value to the height reported by NeuroMorpho.Org. If the values did not match, we computed a conversion factor and applied it to size-related morphometric features including width, height, total length, total surface, total volume, maximum Euclidean distance, and maximum path distance. The specific archives that underwent rescaling and the calculations for the scale correction are detailed in the Supplementary Material at https://github.com/Masood-Akram/Classification_Neurons-Glia/tree/main/Supplementary_Material

When we concluded the main analysis for this work (Fall 2021), a new version of NeuroMorpho.Org had been released (v.8.1). We thus identified an additional balanced dataset of 4292 neurons and 4286 glia, up to version 8.1.90 (December 2021), allowing us to test the robustness of our main results on a completely independent dataset. The scale correction details and the complete metadata breakdown for this additional dataset are also included in the Supplementary Materials at https://github.com/Masood-Akram/Classification_Neurons-Glia/tree/main/Supplementary_Material

### Dimensionality reduction

We computed the coefficient of determination (R^2^) to quantify the pairwise correlation (Di Bucchianico, 2008) among the 19 morphometric parameters across neurons and glia using the *rcorr* function in the R package *Hmisc* (Harrel & Dupont, 2021). We then used Principal Component Analysis (PCA) to reduce the feature redundancy. PCA transforms the data into a set of new orthogonal variables by identifying the directions (principal components) along which the variation in the data is maximal (Abdi & Williams, 2010). By discarding the least informative components, each sample can be represented by fewer, linearly independent features instead of more, mutually dependent variables (Ringnér, 2008). Thus, PCA reduces the dimensionality while retaining most of the variation of the data. PCA was performed with the R package *stats* (Core Team, 2021) by using the function *prcomp()* and setting *scale = TRUE.* Along with PCA, all parameters were standardized by first subtracting the mean of the entire feature vector from each element and then dividing by the standard deviation.

### Supervised learning

Training data consisted of the normalized principal components of the morphometric features as input, and the known class labels (the cell identity ‘neuron’ or ‘glia’) as output. We used three distinct classification algorithms implemented in the R programming language (Core Team, 2021) v.4.1.1 for Windows. In all cases we calculated sensitivity, specificity, and accuracy, respectively defined as the fractions of true positives, true negatives, and correctly classified (true positives plus true negatives) cells, using the *caret* (Kuhn, 2021) package in R (Core Team, 2021).

### K-Nearest Neighbor (KNN)

is a supervised learning algorithm that can be used both for classification (discrete value output), as applied here, and regression (continuous value output) problems. In KNN, the training instances are stored with their labels and each new instance is compared with the labeled ones using a similarity matrix. The vote for each new instance’s label by comparing to existing instances is taken from the value of *k.* For example, if k is set to 5, 5 nearest neighbors are identified from the training instances and the class label with the highest frequency is assigned to the new instance (Aha et al., 1991). The default Euclidean distance was used here to compute similarity between two data points. The built-in *caret* package (Kuhn, 2021) was utilized for the KNN implementation by using the *train()* function and setting *method = “knn”* with *tuneLength = 10* and *k = 5.*

### Support Vector Machine (SVM)

is a binary classification algorithm based on finding the maximum margin hyperplane that gives the greatest separation between the data points of different classes in multidimensional space. Those data points closest to the hyperplane are called the support vectors. If the data are not linearly separable, different kernels can be selected for nonlinear classification. This classifier is robust to large number of variables and small sample sizes (Cortes & Vapnik, 1995). We implemented SVM using the *caret* package (Kuhn, 2021) using the *train()* function with *tuneLength = 10,* and *method = “radial”* kernel, which gave the best classification accuracy and is also a common choice for classification tasks (Luts et al., 2010).

### Random Forest (RF)

consists of a large number of individual decision trees. Each individual tree in the forest splits out a class prediction and the most frequent class becomes the model prediction. This is one of the most popular machine learning algorithms and is capable of both classification (as used here) and regression (Breiman, 2001; Sarica et al., 2017). We applied the *randomForest* package (Liaw & Wiener, 2002) using function *train(), method = “rf”,* and *ntree* = 500. The rationale for this choice is that a relatively high number of decision trees ensures that every input row is predicted multiple times. Parameter *mtry* determines the number of variables randomly sampled as candidates at each split and was set to the default value of 5.

### K-fold Cross Validation (K-fold CV)

It is customary in supervised learning to train the model on the majority of the data, leaving the remaining for testing. To rigorously examine the classification performance on our data, we performed K-fold cross validation. This process divides the dataset into *k* equal parts. A classifier is first trained on *k-1* parts for each fold. The accuracy of the trained model is then assessed by using the part of data excluded from the *k-1* parts in training (Bouckaert, 2003). We performed 10-fold CV repeated 10 times using the *caret* (Kuhn, 2021) package by using the function *trainControl()*, *method = “repeatedcv”*, *number = 10,* and *repeats = 10.*

### PSwarm

is a global optimization solver for bound and linearly constrained problems (Vaz & Vicente, 2009). This algorithm is based on a pattern search and particle swarm method, which guarantees the convergence to stationary points from arbitrary starting points. We used the PSwarm Solver (v.1.5, June 2020, norg.uminho.pt/aivaz/pswarm/) implementation in R to find the linear discriminant of neurons and glia based on two morphometric parameters. We set the lower and upper bounds to 75 and 175, respectively, for intercept and to −10 and 75 for slope, and the number of iterations *(maxit)* to 2·10^9^. All analyses were carried out on a 64-bit machine equipped with an Intel Core i7-8565U and 16 GB of RAM running Windows 10. The R scripts utilized in this work are released open source at https://github.com/Masood-Akram/Classification_Neurons-Glia/tree/main/R_Code.

### Average Branch Euclidean Length (ABEL)

is the average over all branches in a cell of the straightline distance between the beginning and ending points of each branch. This quantity was calculated from three of the morphometric parameters provided for each cell by NeuroMorpho.Org: branch path (geodesic) length, the number of branches, and contraction, which is the ratio between Euclidean branch length and branch path length (its inverse is tortuosity). Specifically, ABEL was derived by summing the product of contraction by branch path (geodesic) length and then dividing the result by the total number of branches in each cell:

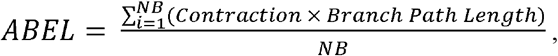

where NB is the total number of branches. We also calculated ABEL of the terminal branches (from a bifurcating point to the tip) and of internal branches (between two consecutive bifurcation points) of both glia and neurons from the .swc reconstruction files provided by NeuroMorpho.Org using L-Measure. In particular, for every cell we first extracted path length and contraction values for each branch while setting “Terminal_Degree=1” under Specificity for terminal branches and “Terminal_Degree>1” for internal branches. We then multiplied the path length and contraction values and took the average within each group (terminal, internal). Lastly, we were also interested to determine the classification power of ABEL when only a small sample of branches was used to estimate the ABEL value. To this aim, we first extracted for every cell path length and contraction values of all branches with L-Measure without setting any Specificity (thus including both internal and terminal branches) and multiplied each pair of values to obtain the Euclidean lengths of all branches. We then utilized the *random* library (Van Rossum, 2020) in Python 3 (Van Rossum & Fred L., 2009) to stochastically select 100 sets of N values without replacement, where N varied from 1 to 15. The N values were used to compute ABEL within each set, and the average and standard deviation were then computed over the 100 sets. Finally, classification was carried out using the mean ABEL value. The code for this analysis is released open source at: https://github.com/Masood-Akram/Classification_Neurons-Glia/tree/main/Python_Code.

### Average Branch Euclidean Length (ABEL)

All morphological reconstructions utilized in this work are available at NeuroMorpho.Org. The individual archive listing, the metadata, and the scaling adjustments are all included in the Supplementary Materials. The source code for all analysis is publicly available on github.com

## Results

The morphological reconstructions of glial processes and neuronal dendrites utilized in this work were contributed to NeuroMorpho.Org by over 250 independent laboratories (listed in Supplementary Materials) and reflect the distribution of published arbor tracings in neuroscience (Fig. 1). Consistent with this multifarious provenance, the dataset spans a broad diversity of experimental methodologies, including over 20 different staining methods (e.g., genetic green fluorescent protein labeling, intracellular biocytin injection, immunostaining, and rapid Golgi), 15 digital reconstruction software (Neurolucida, Imaris, Amira, NeuronJ, Simple Neurite Tracer, Vaa3D, Knossos, NeuTube, etc.), and a continuum of ages across the developmental, from embryo through juvenile to old adults. Moreover, the data came from both mammalian and non-mammalian species and a large variety of anatomical regions but were balanced between neurons and glia across these dimensions (Fig. 2). The full breakdown of all metadata categories annotated in NeuroMorpho.Org is provided in Supplementary Materials.

**Figure 1.**
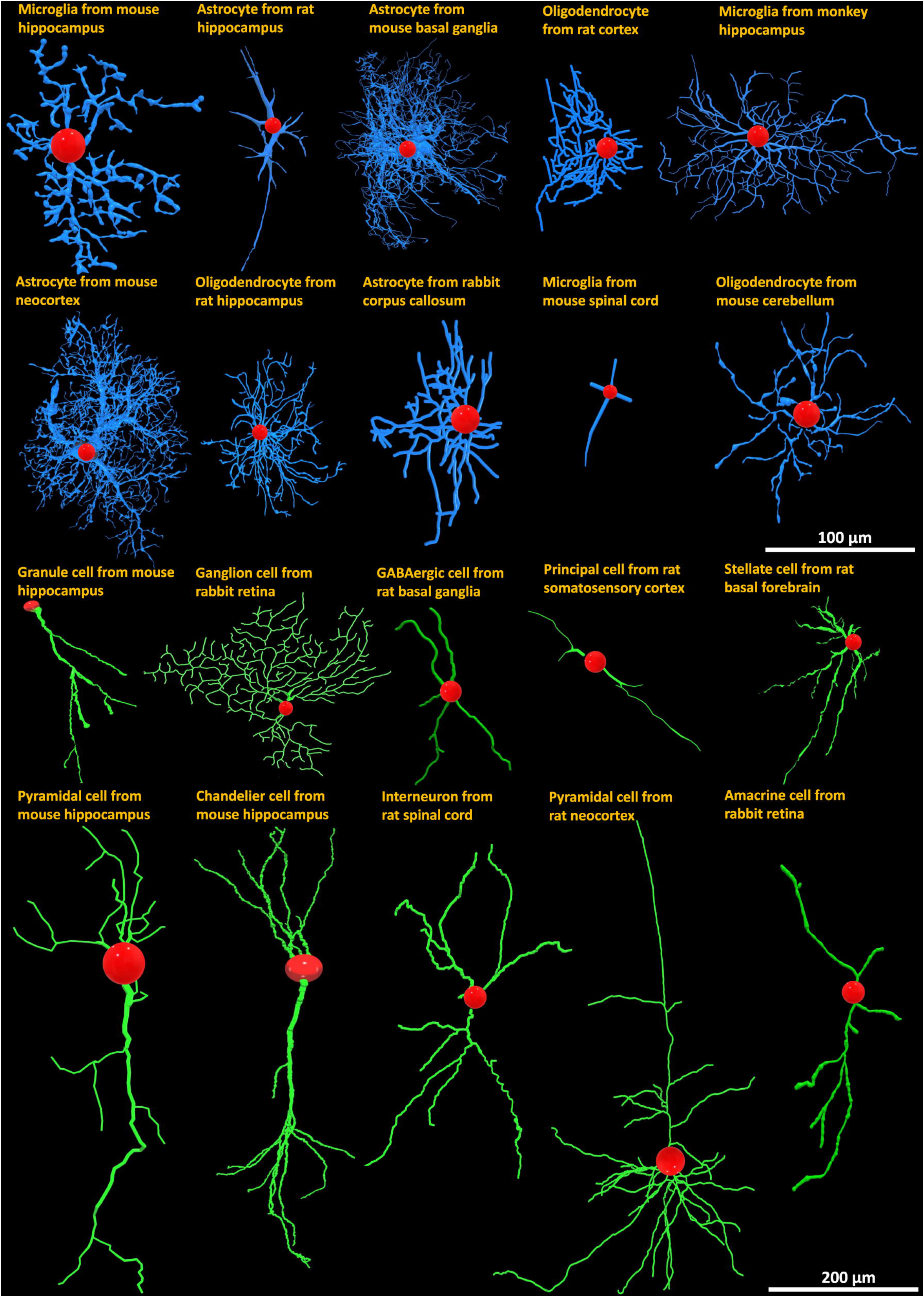
Representative diversity of morphological reconstructions of glia and neurons from NeuroMorpho.Org with labels indicating animal species, anatomical region, and cell type. Blue: glial processes; green: neuronal dendrites; red: cell bodies.

**Figure 2.**
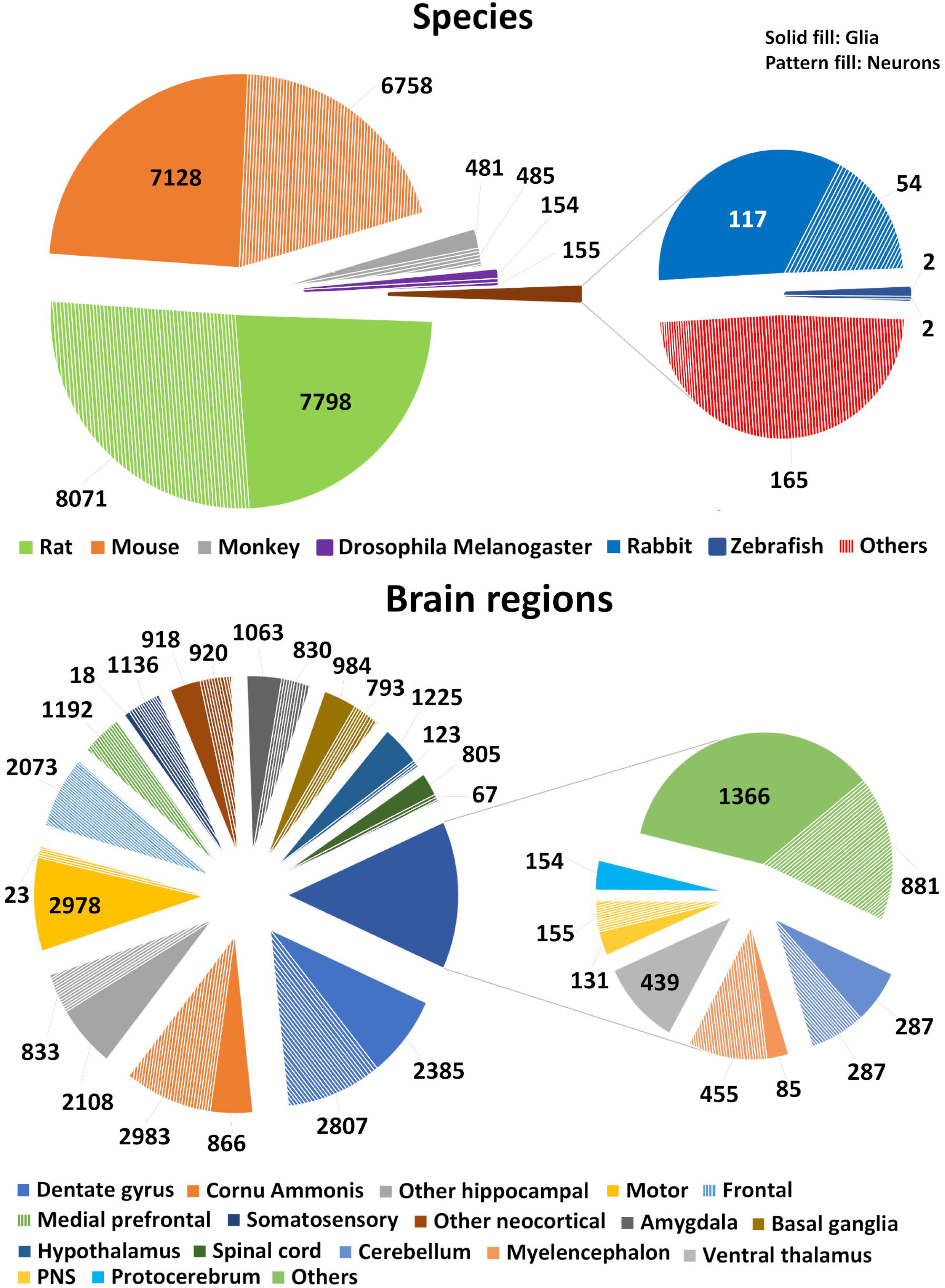
Balanced distribution of (**A**) animal species and (**B**) brain regions for the analyzed glia and neuron datasets.

The morphometric quantification of neural trees supplied by NeuroMorpho.Org provides a detailed 3D representation of branch geometry (Fig. 3). The extracted features include parameters characterizing both the overall size of the arborization and the scale-invariant properties. The formers include total cable length and surface area, spanning height and width, maximum Euclidean and path (geodesic) distance from the root (soma), and average branch diameter among others. The latter measurements capture bush complexity (e.g., number of branches and tree stems), branch angles (local and remote bifurcation amplitude), topological imbalance (partition asymmetry and maximum branch order), and spatial meandering (contraction and fractal dimension), among others. Altogether, this set of morphometric parameters is well suited to characterize the structure of neuronal dendrites and glial processes alike, and thus to quantify their similarities and differences.

**Figure 3.**
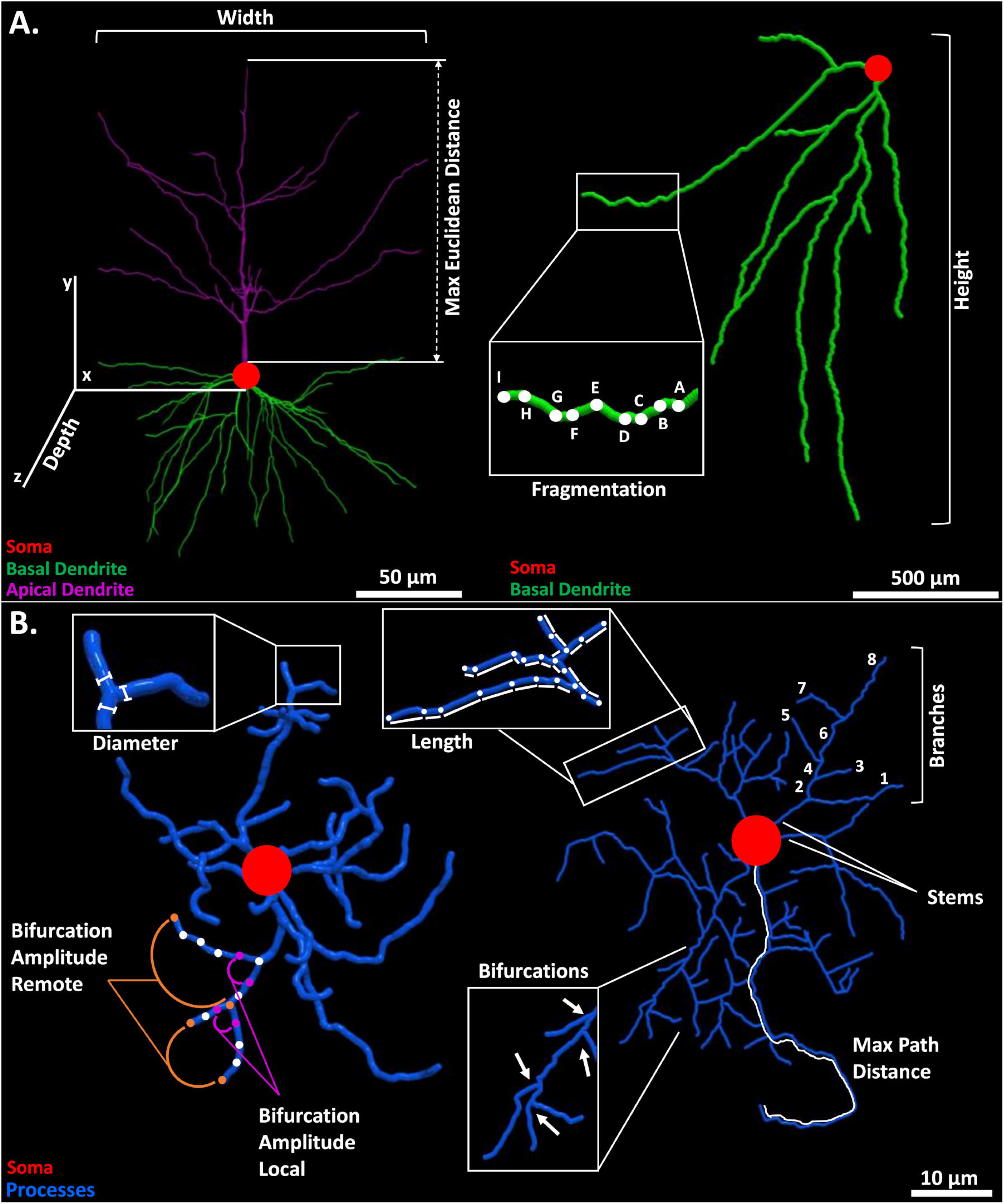
Schematic of selected morphometric features. (**A**) Illustration of width, depth, and maximum Euclidean distance (left) in a monkey neocortical pyramidal cell (NMO_00002) from the Wearne_Hof archive (Duan et al., 2002); and of height and fragmentation (right) in a hippocampal granule cell (NMO_73103) from the Diaz archive (Sebastián-Serrano et al., 2016). (**B**) Diameter and local or remote bifurcation amplitude (left) in a rat neocortical microglia (NMO_95641) from the Roysam archive (Megjhani et al., 2015); and maximum path distance, length, and number of branches, bifurcations, and stems in a rat cortical oligodendrocyte (NMO_131081) from the Sato_Bigbee archive (Mohamed et al., 2020).

Although the above-described parameters are intuitively interpretable, they may not be completely independent of each other. For example, total tree length, surface area, and average branch diameter are expected to be interrelated. This information redundancy can unduly bias the objective characterization of the structural differences between glia and neurons, complicating subsequent interpretations. The pairwise coefficients of determination (R^2^) for glial (Fig. 4A) and neuronal (Fig. 4B) morphometrics confirm the substantial correlation between specific features. For example, surface is highly correlated to volume, the number of bifurcations to the number of branches, length, fragmentation, and branch order (and the latter four to one another), path distance to Euclidean distance, and contraction to fractal dimension. Although the coefficients of determination tended to be higher in neurons than in glia, most visibly between maximum path distance and total surface area, and between overall height and maximum Euclidean distance, the majority of correlations were highly consistent between the two cell types. In order to remove the interdependency among features, we performed PCA jointly on the full dataset to orthogonalize the morphometric parameters (Fig. 4C). The first 11 principal components captured 95.70% of the variance and we thus decided to exclude the last 8 components from machine learning. The 11 principal components considered in subsequent analysis constitute a combined transformation of all 19 morphometric parameters described above but are guaranteed by PCA to be independent.

**Figure 4.**
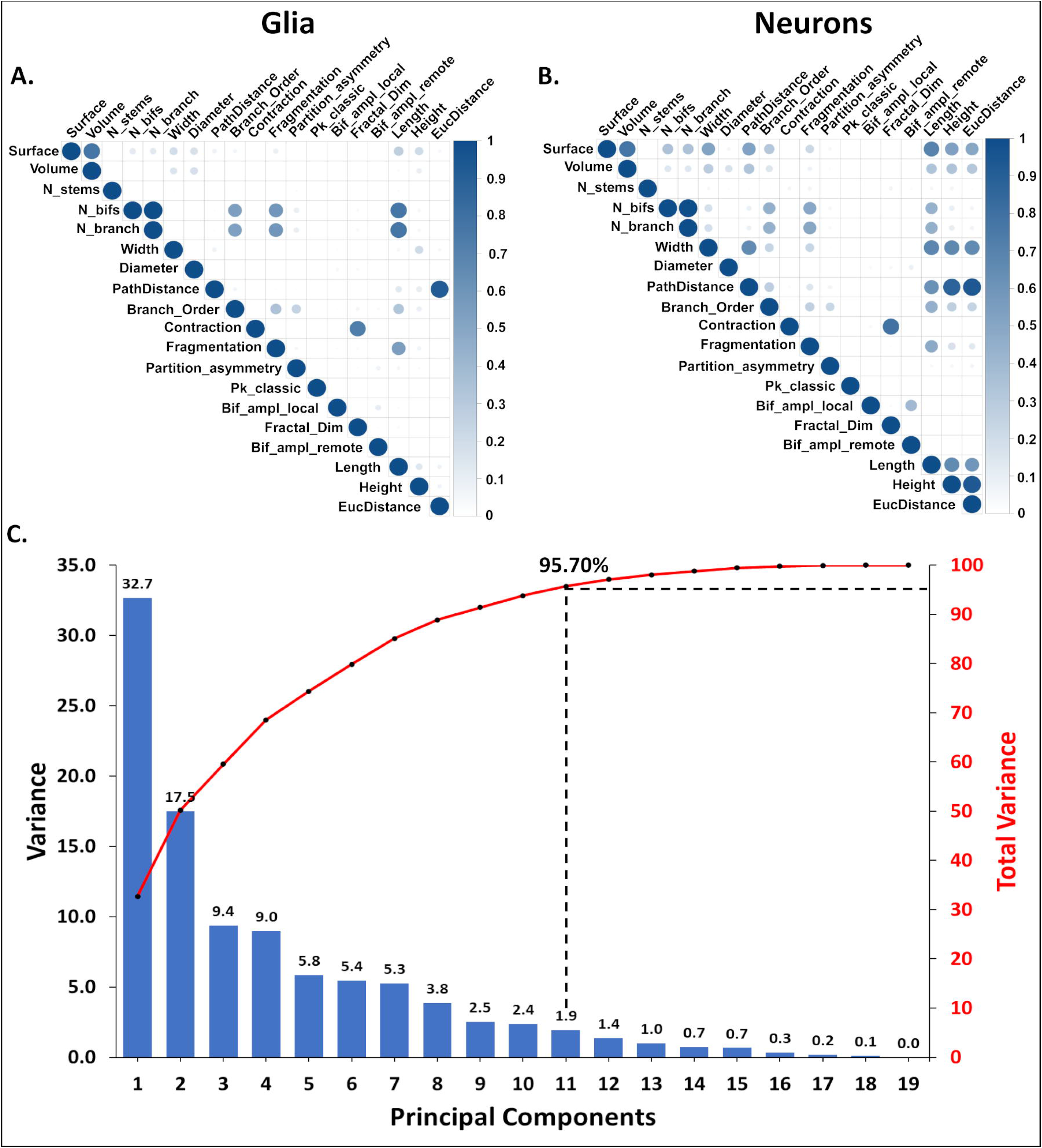
Orthogonalization of morphometric features. (**A**) Correlation matrix quantifying the interdependence among 19 morphometric features of glia and of (**B**) neurons. The coefficient of determination (R^2^) is shown on a dark intensity scale. (**C**) Scree plot of the variance contributed by each sequential principal component (blue bars, left axis) and the corresponding cumulative distribution (red line, right axis).

The first two components (PC1 and PC2) alone capture more than 50% of the overall morphological variance in neural cells. A striking separation between cell classes is apparent on the PC1-PC2 projection plane, with neurons more abundant towards positive coordinates and glia towards negative in both dimensions (Fig 5A). Data points that are close to each other in this projected space represent structurally similar cells, whereas morphologically different cells occupy distant positions. The first two principal components consist of distinct linear combinations of morphometrics: PC1 (Fig. 5B) has strongly positive loading on size (e.g., total cable length, overall arbor height, maximum path distance), while PC2 (Fig. 5C) has strongly negative loadings on tree complexity and other scale-invariant measures, such as number of bifurcations, maximum branch order, and fractal dimension. These results therefore confirm that neurons have greater overall arbor size than glia, as quantifiable by multiple alternative metrics. Furthermore, this analysis reveals that, compared to neuronal dendrites, glial processes tend to form bushier trees, with more symmetric branching distributions and wider bifurcations angles.

**Figure 5.**
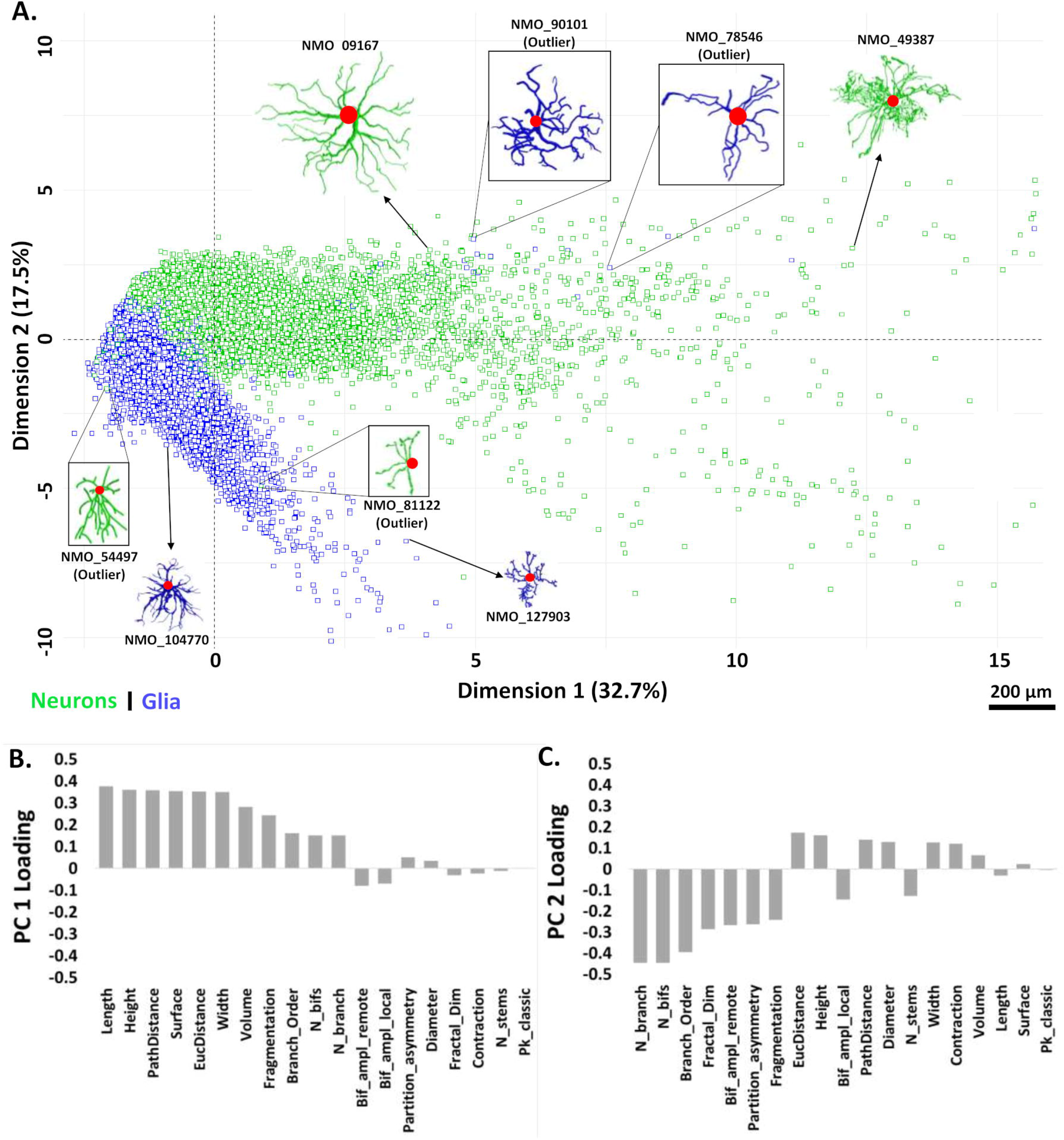
(**A**) PCA biplot of the 2-dimensional distribution of neurons and glia relative to the first two principal components (PC1 and PC2). Morphological tracings of several cells (glia: blue; neurons: green) are also shown to illustrate their structural variability and similarity in this space. (**B**) Linear contributions of all morphometric parameters to PC1 and (**C**) PC2. Negative loadings indicate a high weight of the scale low-end for a parameter: for instance, cells with large positive PC2 values tend to have very few branches, whereas cells with many branches tend to have large negative PC2 values.

The above analysis suggests that neurons and glia may be reliably recognized based on morphological features alone independent of numerous confounds such as species, anatomical region, and experimental methods. To test this hypothesis, we used the 11 principal components explaining >95% of the variance for classification with three supervised learning algorithms: Support Vector Machine (SVM), K-Nearest Neighbors (KNN), and Random Forest (RF). In all cases we performed 10-fold cross validation: the dataset was randomly split into 10 folds without replacement, with 90% of the data used to train the classifier and the remaining 10% used for testing. The process was repeated 10 times for more reliable assessment. The total runtime for 10 repeats of 10-fold cross validation was 15 minutes for KNN with 5 nearest neighbors *(k=5),* 2.2 hours for SVM, and 5 hours for RF. All three classifiers performed remarkably well in separating glia from neurons (Fig. 6). SVM slightly outperformed KNN in terms of sensitivity, specificity, and accuracy, with RF displaying intermediate performance metrics. However, all classification measurements fell within 1% difference for the three algorithms, and the area under curve (AUC), a robust measure of predictive modeling accuracy, was >99.5% for each of them.

**Figure 6.**
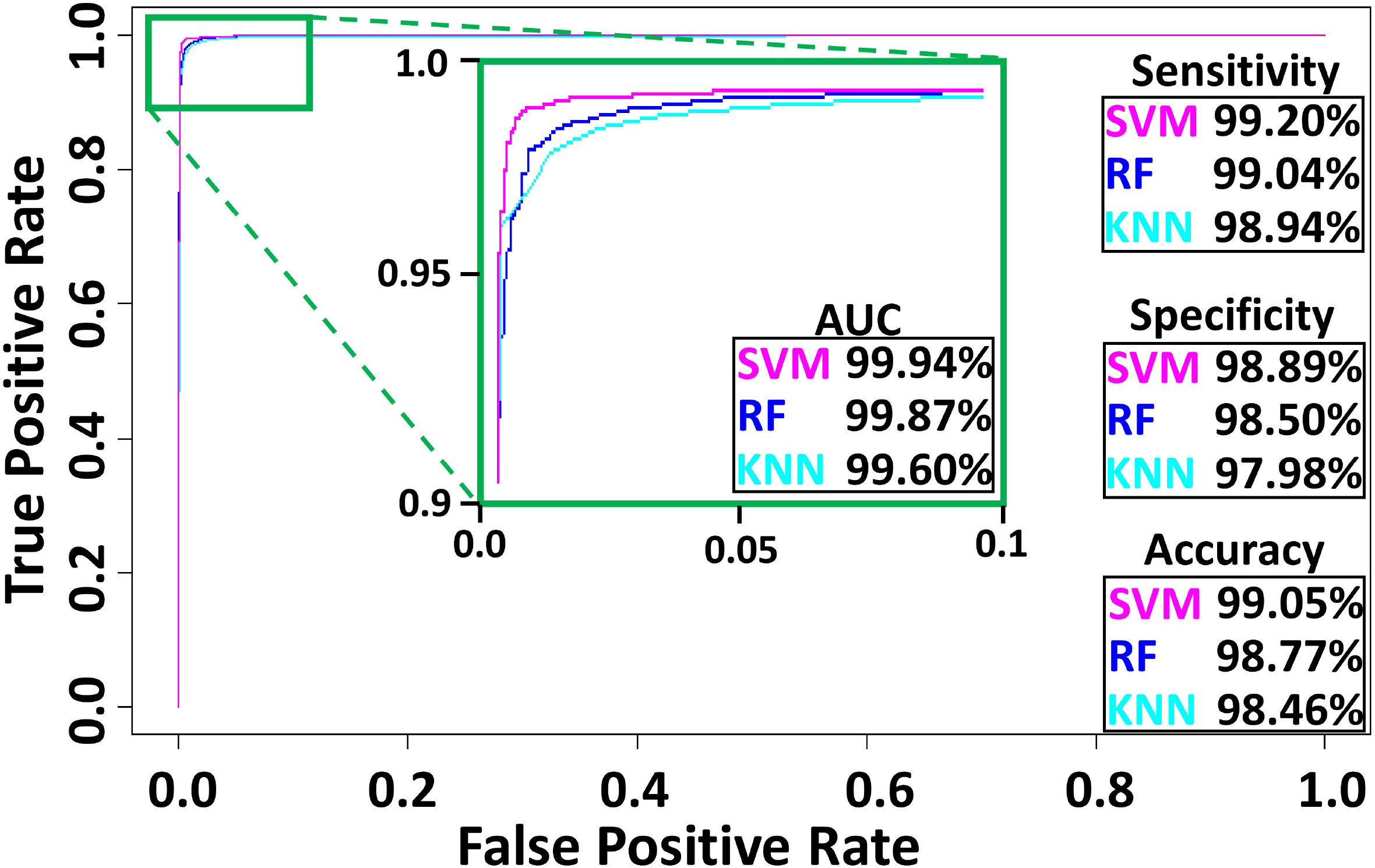
Classification performance for Support Vector Machine (SVM), K-Nearest Neighbors (KNN), and Random Forest (RF), including the area under the curve (AUC) of the Receiver Operating Characteristic plot.

The supervised classification results clearly demonstrate that a variety of automated methods can reliably distinguish glia from neurons by using morphological features alone. Given that the principal components utilized to train the machine learning algorithms represent an extensive battery of morphometric measurements, the question remains whether individual geometric features can be identified that achieve similarly robust performance. Based on the PC1 and PC2 loadings described above, measures of arbor size such as overall height and total length and measures of arbor complexity such as number of branches could constitute viable candidates. Although the corresponding silhouette profiles (Fig. 7) corroborated the expected statistical differences, it also revealed extensive overlap in the corresponding data distributions. For example, the optimal height threshold to discriminate neurons from glia (76.15 μm) resulted in a suboptimal classification accuracy of <0.95, with >6% of neurons misclassified, and even worse performance for total length, number of branches, and any other individual parameter. We reasoned, however, that since neurons have longer cable and glia have more branches, an appropriately combined feature could achieve a multiplicative improvement in discrimination. Specifically, dividing length by number of branches, which defines average branch path length, should yield parameter values with an even larger ratio between neurons and glia than length alone. Moreover, since neurons have slightly less tortuous branches than glia, as indicated by larger (if only marginally) contraction values, multiplying branch length by contraction and averaging over all branches, which defines average branch Euclidean length (ABEL), should further increase corresponding parameter value between neurons and glia. Silhouette analysis confirms the considerably better separation between neurons and glia based on ABEL when compared to all other individual morphometrics (Fig. 7).

**Figure 7.**
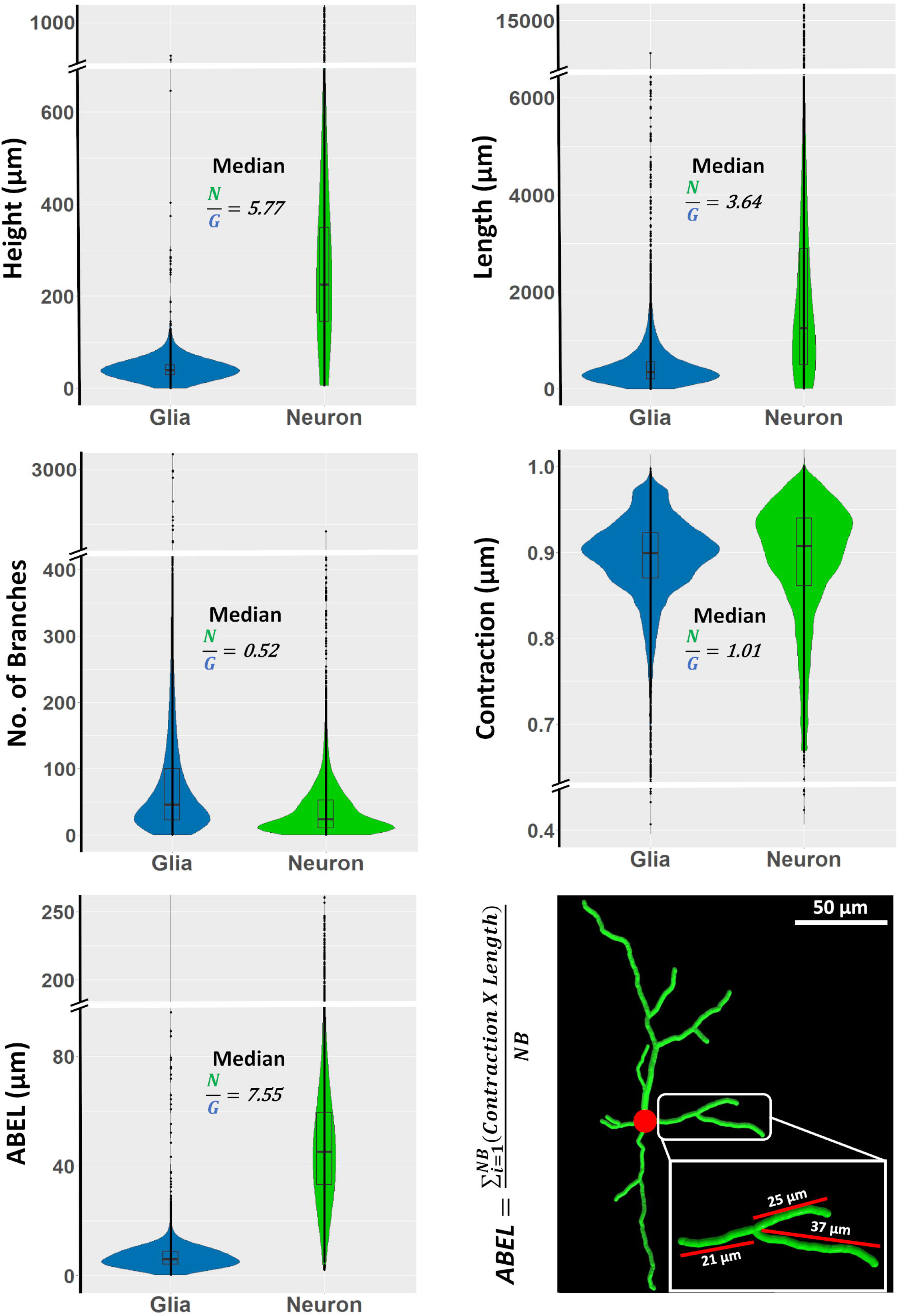
Silhouette profiles of length, height, contraction, number of branches, and average branch Euclidean length (ABEL) of glia and neurons, and examples of branch Euclidean length measurements from a rat basal ganglia GABAergic cell (NMO_68194) from the Smith archive (Smith et al., 2015).

The optimal ABEL threshold of 14.33 μm results in overall classification accuracy of 97.6%, with fewer than 2.4% of glia and 2.5% of neurons misclassified. Notably, the misclassification rate dropped steeply around the threshold ABEL value, with >90% of the misclassified cells found in a narrow ABEL range of 7 (12-19) μm (Fig. 8A). These results were robust across multiple species, strains, developmental stages, anatomical regions, types of glial and neuronal cells, labeling techniques, and experimental methods, as detailed in the Supplementary Materials. For example, when dividing all data by contributing labs, for more than three-quarters of cases the misclassification rate was less than 1%. The rare exceptions consisted of specific phenotypes as discussed at the end of the Results. Furthermore, even an incomplete sampling of neural branches is sufficient for reliable classification based on ABEL: the accuracy is essentially unaltered when using 15 randomly chosen branches (97.3%) and remains above 95% when reducing the ABEL sample size to 5 branches (Fig. 8B). To determine if the classification could be improved further by considering arbor height together with ABEL, we combined the two measures (Fig. 8C). The optimal linear boundary separating neurons and glia followed the equation A=-0.1352H+23.04 μm, where A and H stand for ABEL and height, respectively. This combination increased the classification accuracy of glia and neurons only marginally compared to using ABEL alone, from 97.6% to 98.5%. Altogether, these results indicate that ABEL is an effective, *novel morphological biomarker* for identifying the main neural cell class.

**Figure 8.**
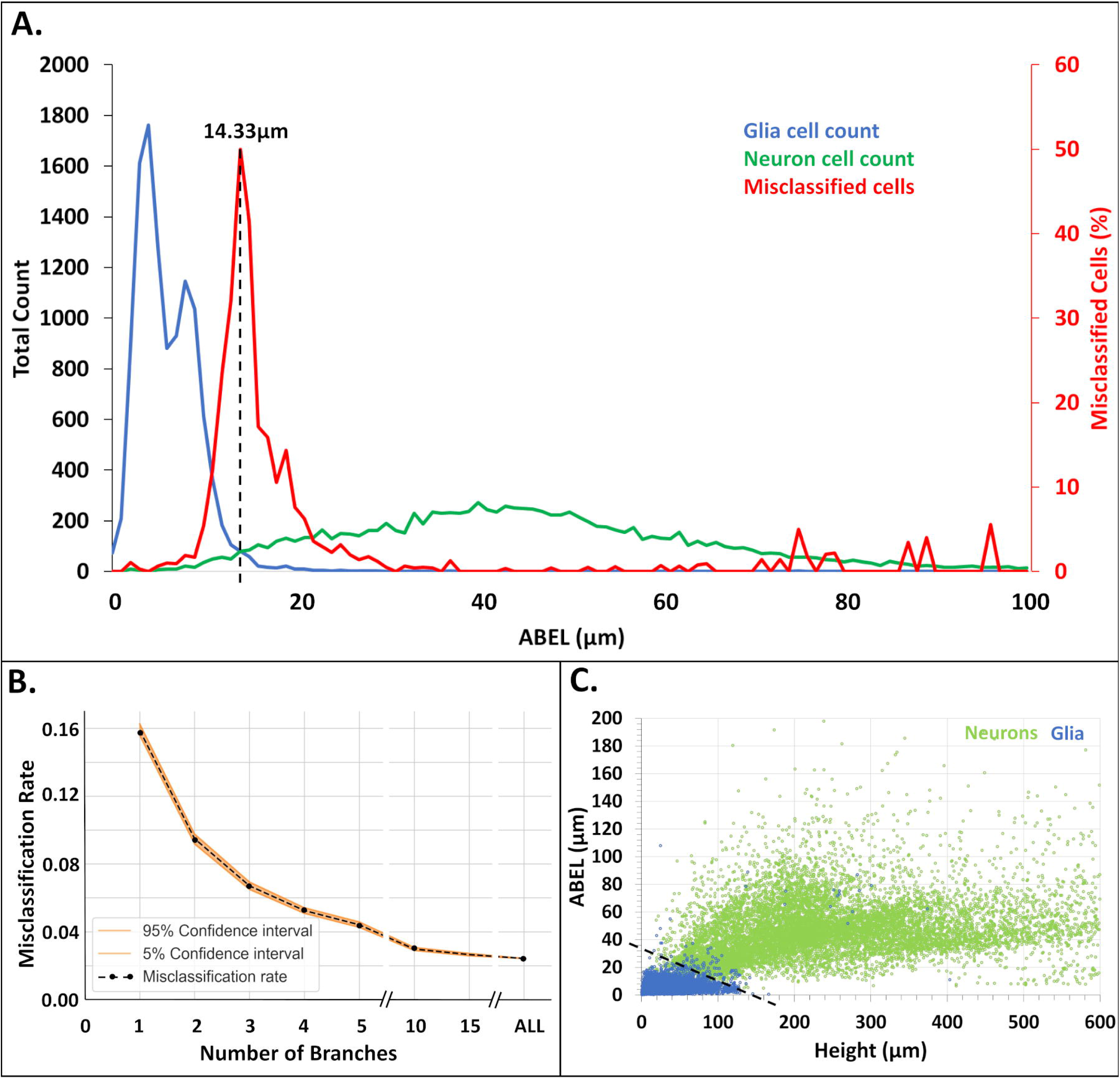
Classification performance of average branch Euclidean length (ABEL). (**A**) ABEL distributions of neurons (green), glia (blue), and cells that are misclassified (red, secondary axis) based on optimal separation threshold of 14.33 μm (vertical dashed line). (**B**) Misclassification rate as a function of the number of branches sampled to estimate ABEL. (**C**) Linear separation (black dashed line) between neurons (green) and glia (blue) on the plane defined by arbor height and ABEL.

Multiple studies reported that certain neuron types have longer terminal branches than internal (bifurcating) branches (Duan et al., 2002; Kawaguchi et al., 2006; Li et al., 2005), but it is unknown whether the same may be true for glia. Since neurons have greater ABEL values than glia, if glial processes have similar length for their terminal and internal processes, then terminal ABEL might be even more effective than overall ABEL to distinguish neurons from glia. To test this possibility, we extracted terminal and internal ABEL for all cells. The distribution of the ratios between terminal and internal ABEL values had an average of approximately 2 for neurons (Fig. 9A), confirming earlier reports that dendrites tend to have longer terminal than internal branches. In contrast, the distribution of the terminal/internal ABEL ratios had an average close to unity for glia, indicating that this phenomenon is limited to neurons. This was also confirmed by linear regression analysis, where the relationship between terminal ABEL and overall ABEL was essentially described by the identity line for glia, but had a slope above unity for neurons (Fig. 9B). Nevertheless, terminal ABEL did not improve the classification accuracy of glia and neurons compared to overall ABEL: in fact, it was slightly decreased to 97.1%, with an optimal separation threshold of 16.20 μm. To investigate why restricting ABEL measurements to terminal branches failed to improve classification performance, we examined the terminal/internal ABEL ratio specifically for the misclassified cells (Fig. 9C). Interestingly, those outlying neurons with exceptionally low ABEL values also displayed similar length between terminal and internal branches. Conversely, outlying glia with exceptionally high ABEL values had longer terminal than internal branches. Linear regression of terminal ABEL versus overall ABEL for the misclassified cells also confirmed that most cells misclassified using overall ABEL are also misclassified using terminal ABEL (Fig. 9D).

**Figure 9.**
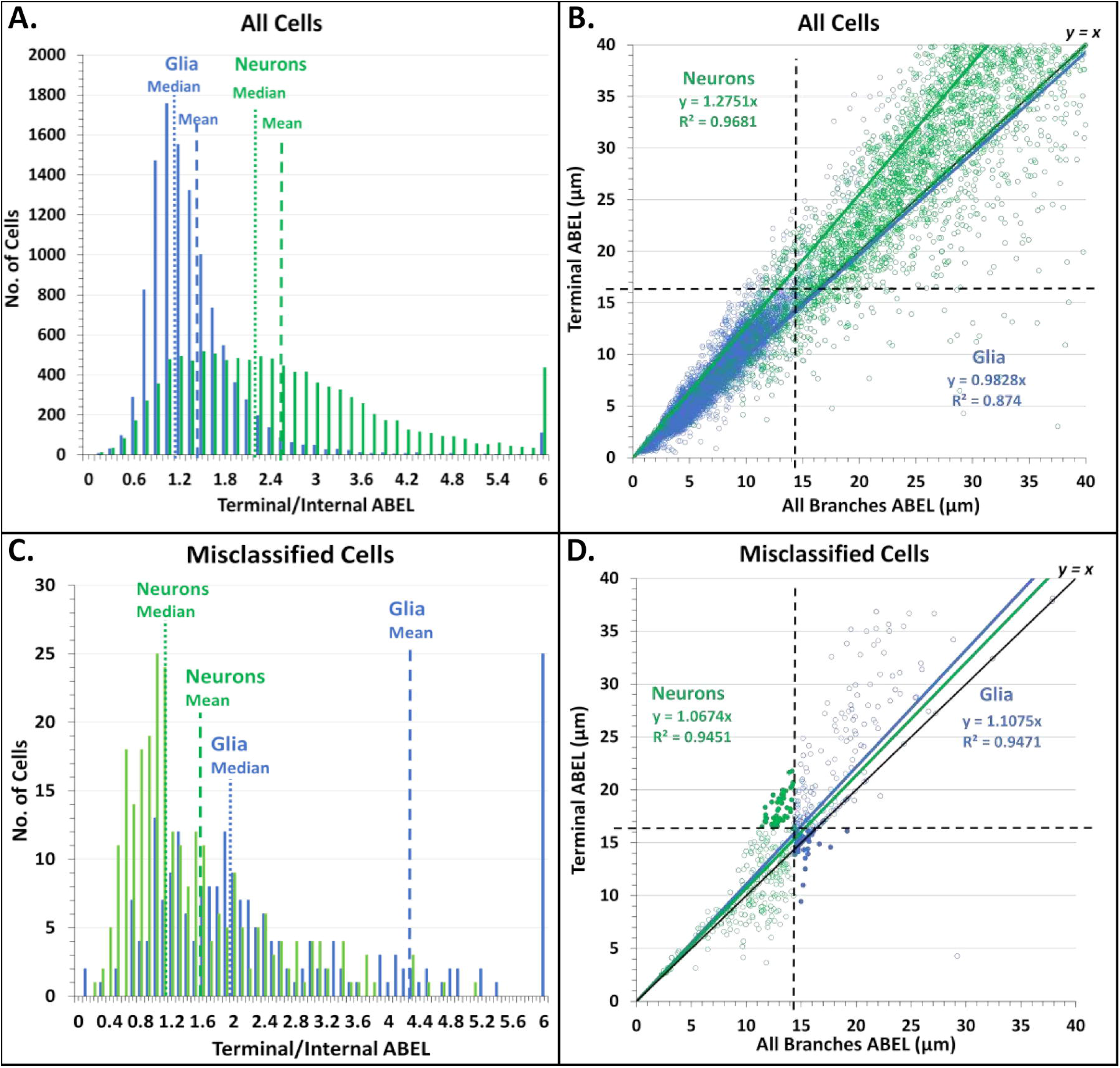
Relationship between the average branch Euclidean length (ABEL) of terminal branches and internal (bifurcating) branches for glia (blue) and neurons (green). (**A**) Distribution of the ratio between terminal and internal ABEL, with medians (vertical dotted lines) and means (vertical dashed lines) indicated. (**B**) 2D scatter and linear regression between terminal ABEL and all-branch ABEL, with respective classification thresholds indicated by horizontal and vertical dashed lines. (**C**) Same as A except limited to cells that are misclassified based on all-branch ABEL. (**D**) Same as B except limited to cells that are misclassified based on all-branch ABEL. The filled circles represent the subset of neurons and glia that are misclassified based on all-branch ABEL but correctly classified based on terminal ABEL. An even larger number of cells (not shown) are correctly classified based on all-branch ABEL but misclassified based on terminal ABEL.

Next, we tested the robustness of ABEL as a morphological biomarker of neurons and glia and how well the optimized classification thresholds generalize to new cell datasets. To this aim, we extended the analysis to the additional glial cells released at NeuroMorpho.Org since the beginning of this study and through the time of this writing (v.8.1.90; N=4,286), balancing the dataset with an equivalent number of neurons (N=4,292) from similar species, anatomical regions, and other metadata (as detailed in Supplementary Materials). The ABEL classification accuracy for this new dataset was the same at 97.6% (using the unaltered 14.33 μm threshold). We then tried to assess whether the few outliers were due to systematic patterns or random noise. Classification accuracy was largely consistent across almost all of the metadata investigated, with only notable exceptions when portioned by brain region (Fig. 10A). Specifically, the high misclassification rate in the cerebellum prompted a deeper evaluation of cells from that region. The misclassified glia consisted of 70 transitional oligodendrocytes and only 1 Iba1-positive microglia, whereas all 78 oligodendrocyte precursor cells, and the rest of cerebellar microglia were correctly classified (Fig. 10B). The 62 misclassified neurons include 8 out of 11 granule cells, and all 54 Purkinje cells, whereas cerebellar basket, stellate, Golgi, Lugaro, and glutamatergic cells were all correctly classified (Fig. 10C). The results indicate that certain cerebellar neurons, specifically Purkinje and granule cells, share similar ABEL with glia. The second, less extreme, exception consisted of the peripheral nervous system (PNS). Here we found that the single culprit was a specific subtype of invertebrate sensory neuron: dendritic arborization (da) Class III cells from the fly larva (46 out of 47 misclassified). In contrast, 104 out of 108 Class I and Class IV sensory neurons, and the quasi totality (98.5%) of PNS glial cells, were correctly classified.

**Figure 10.**
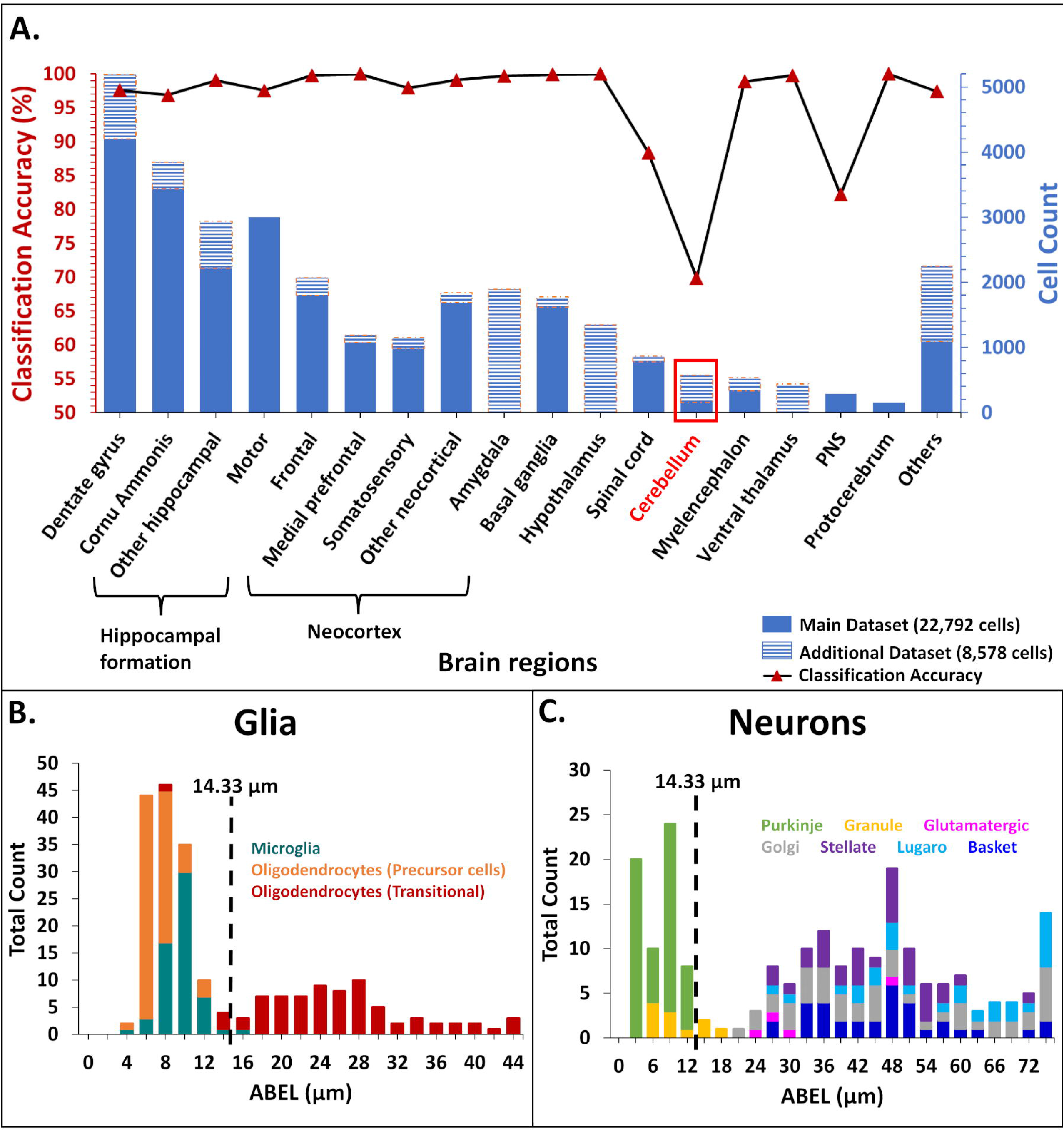
Classification of glia and neurons across anatomical regions. (**A**) Number of cells analyzed (stacked blue bars, right axis: main dataset, solid; and additional dataset, striped) and classification accuracy (black line and red triangle, left axis). (**B**) ABEL distribution of cerebellar glia. Cells to the right of the threshold (vertical dashed line) are misclassified. (**C**) ABEL distribution of cerebellar neurons. Cells to the left of the threshold (vertical dashed line) are misclassified.

## Discussion

Open sharing of digitally reconstructed neuronal morphology from labs across the world has made it possible for researchers to carry out statistical analysis, classification, and computational modeling of their interest (Bota & Swanson, 2007; Halavi et al., 2012; Parekh & Ascoli, 2015). Far fewer morphological classification studies have also included glia, and they typically did not focus on directly comparing neurons to glia. For example, Leyh et al. (2021) classified different types of microglia in healthy and diseased mouse model, while Zhang et al. (2021) added glia as a separate phenotype in a multiclass neuron type categorization task using convolutional neural networks. Recognizing the morphological signatures that distinguish glia from neurons is an important yet unfulfilled step.

This study sought to determine whether neuronal dendrites and glial processes can be reliably separated solely based on their arbor geometries and independent of animal species, anatomical region, developmental stage, and experimental condition. To this aim, we harnessed all publicly available reconstructions of glia and balanced them with an equivalent number of neurons with as closely matching metadata as possible. The resulting dataset of over 30,000 cells spanned the very broad methodological diversity in the field. We then produced a compact, orthogonalized quantification of those morphologies by applying principal component analysis to an extensive battery of extracted morphometrics. Deployment of three traditional supervised learning algorithms yielded exceptionally high (>99%) classification accuracy. We thus set out to determine which specific differences could explain such striking separation. While neurons were confirmed to have larger arbors than glia, we also discovered that glial trees tend to bifurcate more than neurons, and that glial branches are slightly more tortuous than their neuronal counterparts. These features have already proved useful in the separate investigation of neurons (Kawaguchi et al., 2006; Polavaram et al., 2014), and glia (Khakh & Deneen, 2019; Verkhratsky et al., 2019), but to our best knowledge never in their comparison. Combining these measurements, we defined a novel morphometric parameter, the average branch Euclidean length or ABEL, and demonstrated that it constitutes a powerful and robust morphological biomarker of cell type. Throughout the whole dataset, glia had smaller ABEL values than neurons, and fewer than 2.5% of cells were misclassified based on a simple ABEL threshold of ~14 μm. Standard measures of arbor size, such as height, yielded a more than double misclassification rate relative to ABEL.

Molecular expression remains a prominent approach for the consistent identification of cell types in the nervous system. For example, glial fibrillar acidic protein (GFAP), nerve/glial antigen 2 (NG2), and ionized calcium binding adapter molecule 1 (Iba1) are commonly utilized to identify distinct classes of glia. Similarly, neurons are often distinguished by their main neurotransmitter based on presence of vesicular glutamate transporters, glutamic acid decarboxylase, choline acetyl transferase or tyrosine hydrolase. In situ hybridization of the corresponding genes is useful to study the somatic distribution of these neurons and glia but does not reveal their dendrites and processes. Immunolabeling can in some cases visualize cell type-specific neural arbors, and multi-color combinations of antibodies may allow colabeling of distinct cell types in the same preparation. In contrast, relatively simpler but non-selective staining such as Golgi (Ghosh, 2020) impregnates a broad spectrum of neurons and glia. In these cases, ABEL can provide a practical way to quickly recognize neurons from glia. It is important to note in this regard that measuring ABEL does not necessarily require the detailed tracing of the full arbors. Euclidean length is simply defined as the straight-line distance between the start and end points of a branch, which can be computed directly from the microscopic image in any common software. Moreover, we showed that as few as five branches are sufficient to provide an ABEL approximation that distinguishes glia from neurons with >95% accuracy. Even for complex arbors with hundreds of bifurcations and terminations, it is thus possible to estimate ABEL with minimum effort.

Besides the practical utility, it is tempting to speculate about the possible scientific interpretation of our main finding. The systematically small ABEL values of glia suggest a tendency to optimize spatial occupancy, consistent with extensively reported tiling properties for these cells (Barber et al., 2021; Pogodalla et al., 2022). In contrast, the larger ABEL values of neurons are indicative of pressure to maximize spatial exploration, in line with the role of dendrites to integrate converging synaptic signals from multiple neural pathways (Anton-Sanchez et al., 2018; Stepanyants & Chklovskii, 2005). It is especially intriguing to consider the rare exceptions that emerged from our analysis. Since the only glial outliers in terms of ABEL were transitional oligodendrocytes, it is possible that the compact arbor is an acquired property of mature glia rather than an innate feature, and that seeking myelination targets requires a degree of spatial exploration. The main neuronal exceptions were cerebellar granule and Purkinje cells. It may not be a coincidence that these two neuron types together form one of the most peculiar circuits in any neural system: the parallel fibers of the cerebellum, which ascend from granule cell axons and contact the Purkinje dendrites on up to 100,000 spines. Purkinje cells are the output cells of the cerebellar cortex, and their dense, planar dendrites are fan-shaped and branch extensively to cover the field of their respective territories without overlapping (Fujishima et al., 2018). These features, which push Purkinje dendrites towards the compact spatial occupancy typical of glia, are dictated by the need to sample the exceptionally large number of synaptic signals from the parallel fibers (Hirano, 2018). Cerebellar granule cells are the single most abundant neuron type in the mammalian brain (Herculano-Houzel, 2010) as well as the most densely packed (Badura & De Zeeuw, 2017), leading to considerably small dendritic fields (Houston et al., 2017). These characteristics, again determined by the unique connectivity profile of the cerebellar parallel fibers, are more akin to those of glial processes than of typical neuronal dendrites. Of note, the other cerebellar neurons (basket, Lugaro, Golgi, and stellate cells) are all correctly classified by ABEL. These exceptions point to a clearly different cell organization in the cerebellum compared to other brain regions.

Aside from the sparse exceptions, the robustness of the results reported in this study is underscored by the very large dataset, distributed provenance of the reconstructions, and broad diversity of metadata. At the same time, it is also essential to recognize that this study is intrinsically limited by the data availability. For example, although the included species span primates, rodents, fish, and invertebrates, the majority of reconstructions for both neurons and glia come from rats and mice. Furthermore, while many anatomical regions are represented in the study, the list is far from complete. And albeit several classes of glia and of neurons were analyzed, their distribution was far from uniform. These factors reflect the state of the research in this field rather than a flawed analysis design. Nevertheless, the conclusions must be considered tentative until further validated as more data continue to accumulate.

This work also illustrates the usefulness of subjecting very large datasets to exploratory analysis via machine learning, followed by a targeted investigation of the most promising phenomena or patterns revealed. This “breadth-then-depth” approach may help shed light on otherwise elusive mechanisms. In particular, tracing glial morphology has become progressively more common and, thanks to increased awareness of data sharing, ever larger amounts of glial reconstructions are being deposited to NeuroMorpho.Org. This increment in data availability in a public repository opens new doors for scientific discovery, especially when applying different analysis and modeling techniques for glia that have been productively applied to neurons since the early days of computational neuroscience.

## Supplementary Materials

The following files are available at https://github.com/Masood-Akram/Classification_Neurons-Glia/tree/main/Supplementary_Material

Scale Correction Main Dataset: calculations of the correction factors for the archives of the main dataset reporting reconstruction coordinates in pixels rather than microns.

Scale Correction Additional Dataset: calculations of the correction factors for the archives of the additional dataset reporting reconstruction coordinates in pixels rather than microns.

Metadata Dimensions Main Analysis: detailed breakdown of the metadata for all archives of the main dataset.

Metadata Dimensions Additional Analysis: detailed breakdown of the metadata for all archives of the additional dataset.

## Acknowledgments

This work was supported in part by NIH grants R01NS39600, U01MH114829, and R01NS086082 to GAA, and R01EY029715 to QW. We are thankful to the NeuroMorpho.Org team and all data providers whose generous resource sharing made this study possible.

